# MCMC-CE: A Novel and Efficient Algorithm for Estimating Small Right-Tail Probabilities of Quadratic Forms with Applications in Genomics

**DOI:** 10.1101/2025.03.16.643492

**Authors:** Vy Q. Ong, Bich N. Choi, Devin P. Lundy, Hongyan Xu, Santu Ghosh, Hui Jiang, Yang Shi

**Affiliations:** Biostatistics and Bioinformatics Core, Karmanos Cancer Institute and Department of Oncology, Wayne State University School of Medicine, Detroit, MI 48201, USA; Department of Biostatistics, Data Science and Epidemiology, School of Public Health, Augusta University, Augusta, GA 30912, USA; McGovern Medical School at the University of Texas Health Science Center at Houston, Houston, TX 77030, USA; Department of Biostatistics, Center for Computational Medicine and Bioinformatics and University of Michigan Rogel Cancer Center, University of Michigan, Ann Arbor, Michigan 48109, USA

**Keywords:** quadratic forms, *p*-values, cross-entropy method, Markov chain Monte Carlo, importance sampling, genomics

## Abstract

Quadratic forms of multivariate normal variables play a critical role in statistical applications, particularly in genomics and bioinformatics. However, accurately computing small right-tail probabilities (*p*-values) for large-scale quadratic forms is computationally challenging due to the intractability of their probability distributions, as well as significant numerical constraints and computational burdens. To address these problems, we propose MCMC-CE, an innovative algorithm that integrates Markov Chain Monte Carlo (MCMC) sampling with the cross-entropy (CE) method, coupled with leading-eigenvalue extraction and Satterthwaite-type approximation techniques. Our approach efficiently estimates small *p*-values for quadratic forms with their ranks exceeding 10,000. Through extensive simulation studies and real-world applications in genomics, including genome-wide association studies and pathway enrichment analyses, our method demonstrates advantageous numerical accuracy and computational reliability compared to existing approaches such as Davies’, Imhof’s, Farebrother’s, Liu-Tang-Zhang’s and saddlepoint approximation methods. MCMC-CE provides a robust and scalable solution for accurately computing small *p*-values for quadratic forms, facilitating more precise statistical inference in large-scale genomic studies.

## 1. INTRODUCTION

The problem addressed in this work is the computation of the right-tail probabilities (referred to as ***p*-values**) of the quadratic forms of multivariate normal variables (referred to as **quadratic forms**),

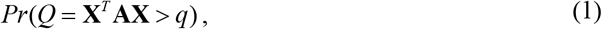

where *q* is a given scalar, **X** = (*X*,…, *X*_*p*_)^*T*^ is a *p*-dimensional multivariate normal random vector following *N*_*p*_ (**μ, Σ**) with mean **μ** = (*μ*_1_,…, *μ* _*p*_)^*T*^ and a *p* × *p* positive definite covariance matrix **Σ**, and **A** is a *p* × *p* symmetric and non-negative definite matrix (Duchesne and De Micheaux, 2010). The dimension and rank of matrix **A** is also referred to as the dimension and rank of the quadratic form *Q*. The form of *Q* can also be written as a weighted sum of independent noncentral chi-squared variables, and in the simplest case where **A** = **Σ** = **I** _*p*_, with **I** _*p*_ the *p* × *p* identity matrix, *Q* reduces to a single noncentral chi-squared variable with *p* degrees of freedom (DFs) and noncentrality parameter (NCP) *δ* = **μ**^*T*^ **μ** ((Duchesne and De Micheaux, 2010); see also the Methods section). Hence, *Q* is also known as the sum (or linear combination) of chi-squared variables (Davies, 1980) or the generalized chi-squared variable (Richter and Schumacher, 2000). The distribution of this quadratic form has broad applications in many statistical hypothesis testing problems (e.g. see (Mathai and Provost, 1992) for book-length discussions), and accurately evaluating its small *p*-values as defined in (1) is of interest in various fields, including high energy physics (Bausch, 2013), signal processing (Hammarwall et al., 2008), wireless communication systems (Issaid et al., 2020), genomics and bioinformatics (Goeman et al., 2004, Liu et al., 2007, Pan et al., 2014, Qu et al., 2013, Wu et al., 2010, Wu et al., 2011). In genomics and bioinformatics, quadratic forms have been used to jointly test the association of an outcome variable with a group of genomics features, such as a group of genetic variants (e.g. single nucleotide polymorphisms, or SNPs) or a group of genes (e.g. a pathway consisting of multiple genes). The former includes methods such as the sequence kernel association test (**SKAT**) (Wu et al., 2011), the sum of powered score (**SPU**) test when the power equals two (Pan et al., 2014), and the linear score test for variance components in linear mixed models (Qu et al., 2013). The latter includes the score test in linear kernel machine regression (Liu et al., 2007) and the score test based on generalized linear models (named **globaltest**) (Goeman et al., 2004). For all these applications in genomics, an accurate estimation of small *p*-values associated with the quadratic form test statistics is essential for multiple hypothesis testing correction and the ranking of genomic features by their statistical significance. In modern genomic studies, extremely small *p*-values, sometimes on the order of magnitude of 10^−50^ – 10^−100^, are not rarely reported (Bangalore et al., 2009).

Most existing methods for computing small *p*-values of quadratic forms have been reviewed and evaluated by others (Bausch, 2013, Chen and Lumley, 2019) and are also assessed and compared with our proposed method in the Results section. Here, we briefly summarize them. One class of methods approximates the distribution of *Q* using well-known distributions through moment-matching techniques, which can be traced back to the famous Satterthwaite approximation (also known as the Welch–Satterthwaite equation) that approximates the sum of central chi-squared variables, a special form of *Q* (see the Methods section for details), by a chi-squared distribution (Satterthwaite, 1946). Other examples include approximations using scaled chi-squared (Goeman et al., 2004) and non-central chi-squared (Liu et al., 2009) distributions.

While these methods are generally straightforward to implement and computationally efficient, their accuracy dramatically deteriorates for *p*-values smaller than 10^−4^ – 10^−5^ (e.g. see (Chen and Lumley, 2019) and our simulation studies in the Result section). Another class of methods, known as exact methods, includes Imhof’s (Imhof, 1961), Davies’ (Davies, 1980) and Farebrother’s (Farebrother, 1990, Farebrother, 1984) methods, which could theoretically achieve arbitrary accuracy for computing the *p*-values of quadratic forms if arbitrary precision arithmetic were available. However, this cannot be achieved in practice (e.g. the machine epsilon, also known as machine precision, for most current computer systems that use IEEE 754 double-precision floating-point arithmetic is 2.22×10^−16^). Consequently, these methods fail for small *p*-values due to numerical limitations and typically break down near machine precision, around 10^−15^ or even 10^−14^ on most current computer systems ((Chen and Lumley, 2019); see also the Results section). Another approach, the saddlepoint approximation (Kuonen, 1999), is accurate for computing small values when *Q* takes the form as the sum of central chi-squares (i.e. when **μ** = (0,…, 0)^*T*^ in (1)), but it is unreliable when *Q* takes the general form as the sum of noncentral chi-squares ((Chen and Lumley, 2019); see also the Results section). Additionally, all these methods require that all eigenvalues of matrix **A** are given. When the rank of **A** is large (e.g. from 1,000 to 10,000, which is common in genomics nowadays due to large sample sizes and numerous genomic features), computing all its eigenvalues, whose time complexity scales as *O*(*p*^3^), becomes very inefficient (Lumley et al., 2018). Furthermore, for a very large matrix **A**, computing and storing **A** itself together with all its eigenvalues requires excessive memory and processing power, further limiting the feasibility of these methods in large-scale genomic applications.

This work aims to address the challenges discussed above, and it is an extension of the work of (Shi et al., 2019), where a general algorithm, MCMC-CE, is developed for estimating small *p*-values associated with a broad class of complicated test statistics using the cross-entropy (CE) method (Chan and Kroese, 2012, Rubinstein, 1999) and Markov chain Monte Carlo (MCMC) sampling techniques. The contributions of this work can be summarized as follows:

### (a) Efficient estimation for small quadratic forms

We develop a practical implementation of the MCMC-CE algorithm for estimating small *p*-values of quadratic forms. Through simulation studies, our algorithm demonstrates superior accuracy and reliability compared to existing methods for small quadratic forms with ranks below 200 (referred to as **small quadratic forms**).

### (b) Scalability for large quadratic forms

For **large quadratic forms** with ranks exceeding 250 and up to 10,000, the original version of MCMC-CE (Shi et al., 2019) also faces the above computational burdens, becoming very slow or even failing to converge. Motivated by a recent work (Lumley et al., 2018), we address the computational bottlenecks for large quadratic forms by integrating leading eigenvalue extraction and Satterthwaite-type approximation, enabling MCMC-CE to estimate small *p*-values of large quadratic forms efficiently. Our approach provides broader applicability than previous methods (e.g. (Lumley et al., 2018)) and achieves better numerical accuracy across diverse settings.

The remainder of this paper is organized as follows. The Methods section details our proposed approaches. The Results section presents extensive simulation studies under various settings with different forms of quadratic forms, comparing the numerical performance of our approach to existing methods, followed by two real-world applications in genomics, a genome-wide association study (GWAS) and a pathway enrichment analysis, where our approach is applied to test the association between genomic features and an outcome using the quadratic forms.

## 2. METHODS

### 2.1. MCMC-CE Algorithm for Small Quadratic Forms

Following (Duchesne and De Micheaux, 2010), the quadratic form *Q* in (1) can be written as a linear combination of noncentral chi-squared variables using the eigenvalue decomposition of **A**,

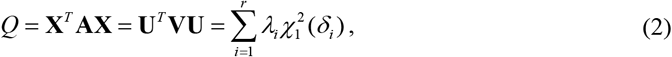

where **V** = diag(*λ*_1_,…, *λ*_*p*_) is a diagonal matrix containing all the *p* eigenvalues of **A**. Some of these eigenvalues can be 0’s, and all the *p* eigenvalues are assumed to be ordered as *λ*_1_ ≥ *λ*_2_ ≥ … ≥ *λ*_*r*_ > 0 and *λ*_*r* +1_ = *λ*_*r* +2_ … = *λ*_*p*_ = 0, where *r* is the number of positive eigenvalues as well as the rank of **A**. Additionally, 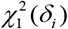 is a noncentral chi-squared variable with 1 DF and NCP *δ*_*i*_, and **U** = **P**(**C**^*T*^)^−1^ **X**, where the matrix **C** corresponds to the Cholesky decomposition of **Σ** satisfying **Σ= C**^*T*^ **C** and the matrix **P** satisfies **PP**^*T*^ **= I**_*p*_ and diagonalizes **CAC**^*T*^, that is, **PCAC**^*T*^ **P**^*T*^ = **V** = diag(*λ*_1_,…, *λ*_*p*_). Thus the distribution of **U** is *N*_*p*_ (**τ, I** _*p*_), where **τ** = **P**(**C**^*T*^)^−1^**μ** and 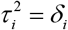, with *τ*_*i*_ the *i*th component of the vector **τ**. Since the zero eigenvalues, *λ*_*r* +1_,…, *λ*_*p*_, do not contribute to the sum 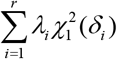, the quadratic form *Q* in (2) can be further simplified as

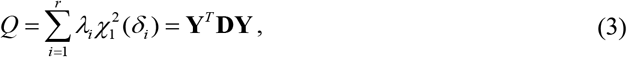

where **D** = diag(*λ*_1_,…, *λ*_*r*_) contains only the positive eigenvalues of **A**, and **Y** follows *N*_*r*_ (**ν, I**_*r*_), with the vector **ν** containing the first *r* elements of **τ**. Due to its simplicity, we will work with the forms **Y**^*T*^ **DY** and 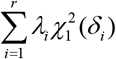 in (3) below.

Given the form of *Q* in (3), the MCMC-CE algorithm, which is the Main Algorithm in (Shi et al., 2019), can be directly applied to estimate the small *p*-value associated with *Q*. By choosing the proposal density *f* (·;**θ**) as a multivariate normal 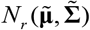, where 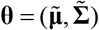 are the CE-optimal parameters that need to be computed in the MCMC-CE algorithm, the following procedure for estimating small *p* = *Pr*(*Q* = **Y**^*T*^ **DY** > *q*) is proposed.

#### Algorithm 1 MCMC-CE algorithm for *p*-values of small quadratic forms.

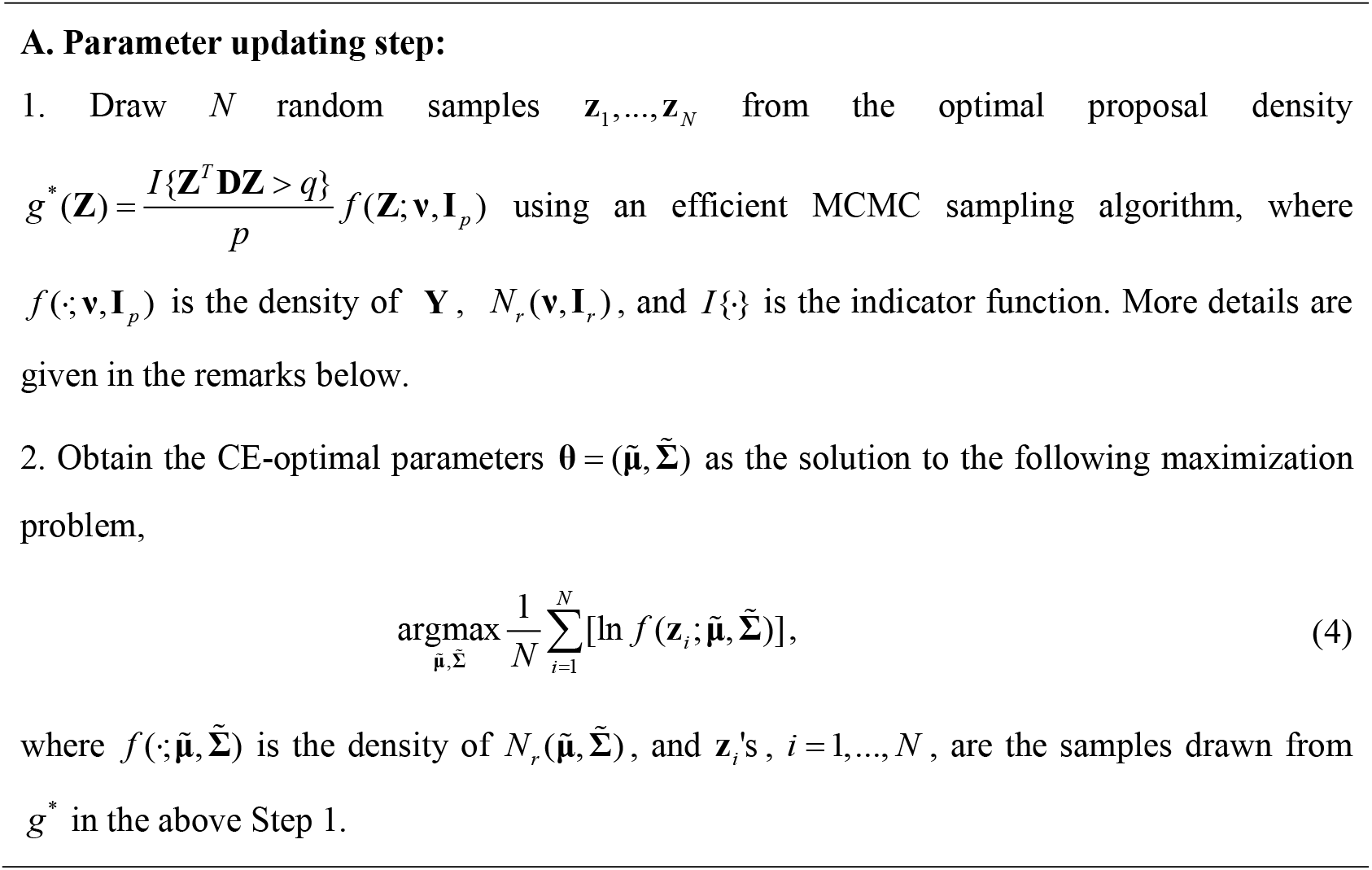
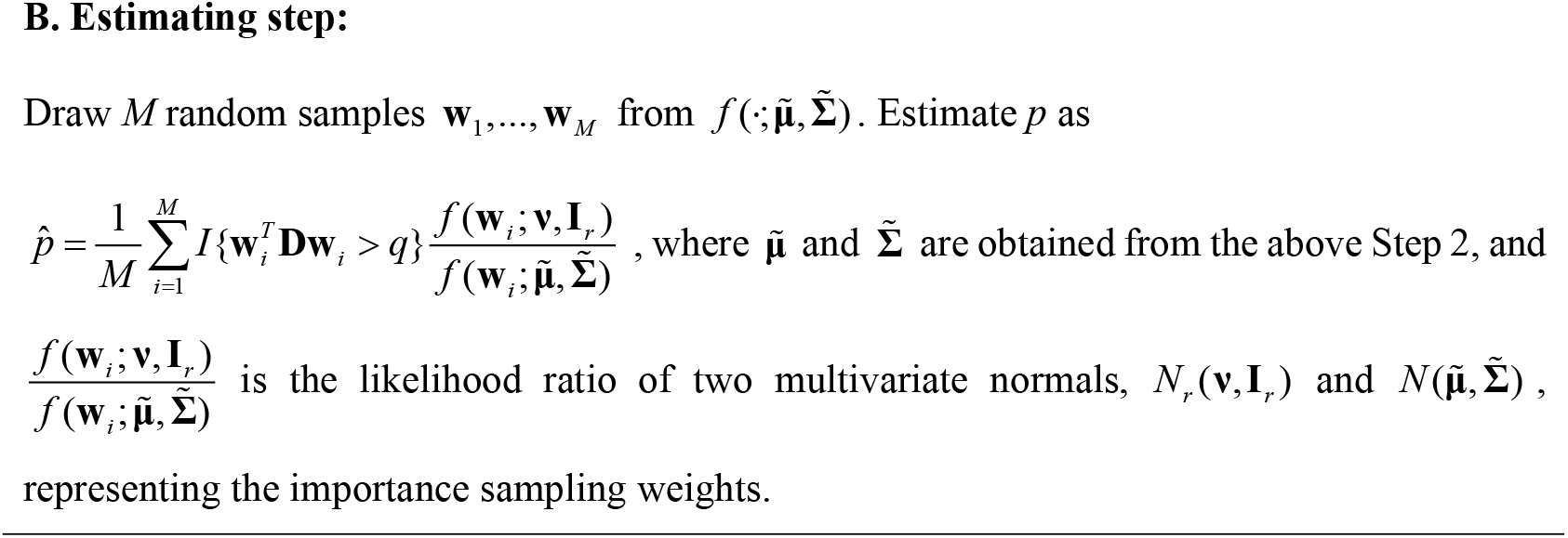

#### Remarks for Algorithm 1

(a) Note that *g*^*^(**Z**) is a truncated probability distribution with the density of **Y**, *N*_*r*_ (**ν, I**_*r*_), mrestricted by the quadratic inequality **Y**^*T*^ **DY** > *q*. There are several MCMC algorithms developed for sampling from such a truncated multivariate normal distribution (Brubaker et al., 2012, Chen and Schmeiser, 1993, Geweke, 1991, Lan et al., 2014, Pakman and Paninski, 2014). Among them, the efficient Hamiltonian Monte Carlo sampler (Pakman and Paninski, 2014) is used in Algorithm 1.

(b) The maximization problem (4) has a closed-form solution, where the solutions to 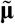 and 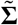 are respectively the sample mean vector and covariance matrix of the drawn samples?**z**_*i*_ ‘s, *i* = 1,…, *N*.

### 2.2. Leading Eigenvalue Approximation for Large Quadratic Forms

When *r*, the rank of matrix **A** is large (e.g. over 250 to around 10,000), Algorithm 1 becomes slow or even fails to converge. Motivated by (Lumley et al., 2018), we propose a scheme that integrates leading eigenvalue extraction and Satterthwaite-type approximation to approximate the distributions of large quadratic forms, enabling MCMC-CE to estimate small *p*-values of large quadratic forms. Specifically, our scheme requires the large quadratic form *Q* to take the following form,

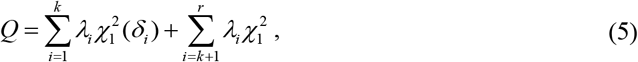

where *λ*_*i*_, *i* = 1,…, *r*, is the *i*th ranked positive eigenvalue of matrix **A** defined in (2). The form (5) requires the NCPs *δ*_*i*_’s, *i* = *k* +1,…, *r*, in the last (*r* − *k*) terms to be 0’s or numerically very close to 0’s, and thus they can be ignored. That is, the first *k* terms in (5) can be either noncentral or central chi-squares, while the last (*r* − *k*) terms need to be central chi-squares. The two sums 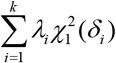 and 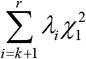 in (5) are referred to as the sum of the *k* leading terms and the sum of the remainder (*r* − *k*) terms, respectively. We utilize a Satterthwaite-type approximation to approximate the sum of the remainder terms, 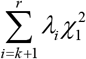, by a scaled noncentral chi-squared random variable, 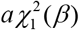, with scale parameter *α*, NCP *β* and 1 DF. Thus, the distribution of *Q* in (5) is approximated using the distribution of

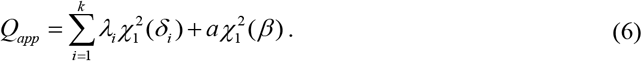

Matching the moments between 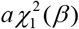 and 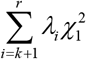 gives

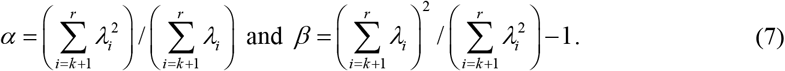

Algorithm 1 can be used to estimate the small *p*-value associated with *Q*_*app*_, which can be seen by writing *Q*_*app*_ in (6) as

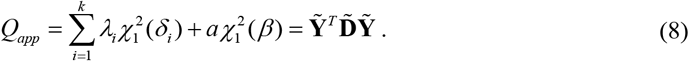

where 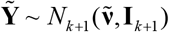, a multivariate normal with 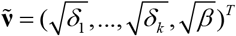, and 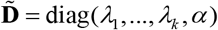. Based on the form of *Q*_*app*_ in (8), the algorithm for estimating 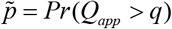 as an approximation to *Pr*(*Q* > *q*) using Algorithm 1 can be summarized as follows.

#### Algorithm 2 MCMC-CE algorithm for *p*-value with large quadratic forms.

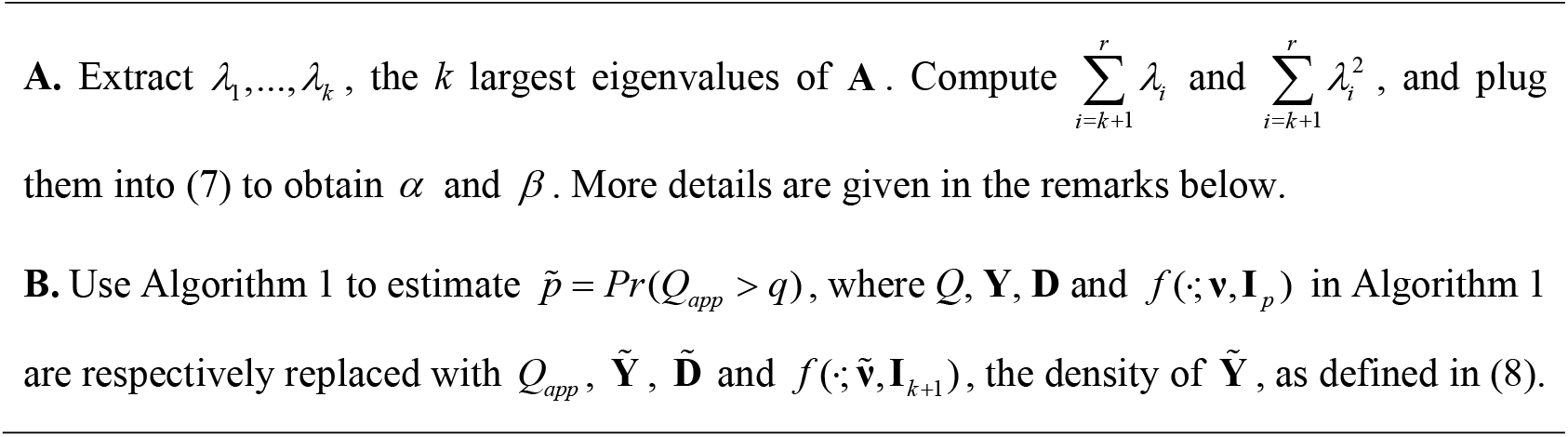

#### Remarks for the leading eigenvalue approximation scheme and Algorithm 2

a. The difference between our approximation scheme and the one in (Lumley et al., 2018) can be summarized as follows. The method in (Lumley et al., 2018) requires the NCPs *δ*_*i*_’s, *i* = 1,…, *r*, in all the *r* terms in (2) to be 0’s, meaning that *Q* must be the sum of *r* central chi-squared variables. Thus, the form of *Q* studied in (Lumley et al., 2018) is a special case of the form (5), whose distribution our scheme approximates. Therefore, our approach has broader generalizations. Taking SKAT (Wu et al., 2011) and *globaltest* (Goeman et al., 2004) as examples, the form of *Q* as the sum of central chi-squared variables corresponds to the distributions of these test statistics under the null hypothesis of no association between the genomic features and the outcome, while the more general form *Q* in (5) corresponds to their distributions under the alternative hypothesis that at least one of the *k* genomic features is associated with the outcome. Thus, our approach not only can estimate *p*-values under the null hypothesis of SKAT and *globaltest* but also can facilitate power analysis under the alternative hypothesis for those tests.
b. Choosing *k*, the number of leading eigenvalues used in Algorithm 2. We find that the empirical scheme proposed in (Lumley et al., 2018) works well in practice, which starts with a reasonable *k* and increases it until the estimated *p*-value stabilizes. Based on (Lumley et al., 2018) and our findings, *k* from 100 to 150 generally provides satisfactory accuracy when the rank of **A** is from 5,000 to 10,000 (see the Simulation Studies). Hence our recommendation is consistent with (Lumley et al., 2018): check if *k* = 100, 150 and 200 give similar estimates. If so, *k* is sufficient; otherwise, *k* needs to be increased.
c. Computing the *k* leading eigenvalues, *λ*_1_,…, *λ*_*k*_, in Algorithm 2. As proposed and studied in (Lumley et al., 2018), at least two approaches can be used: the thick-restart Lanczos method (trLanczos) (Korobeynikov, Wu and Simon, 2000, Yamazaki et al., 2010) and the stochastic singular value decomposition (SSVD) algorithm (Halko et al., 2011). Both trLanczos and SSVD are accurate for large matrices, but SSVD is not accurate for small matrices (Lumley et al., 2018). Their computational speeds are similar (Lumley et al., 2018), and a comparison of their runtimes is presented in the Results section.
d. Computing 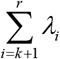 and 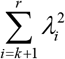 in Algorithm 2. Two approaches are proposed in (Lumley et al., 2018): the fast trace estimator (Hutchinson, 1989) and the stabilizing ratio estimator (Fuller, 2011). Both methods can be used in Algorithm 2, but the fast trace estimator is generally more efficient without significant loss of accuracy (Lumley et al., 2018), making it the default choice.

## 3. RESULTS

We first conduct simulations under various settings to study the numerical accuracy of Algorithms 1 and 2, and compare them with existing approaches. Then we demonstrate the applications of our proposed algorithms on two real-world datasets in genomics, a GWAS dataset and a pathway enrichment analysis, in which small *p*-values of the quadratic forms for testing the association between genomic features and an outcome are estimated. All analyses are performed on a Windows desktop PC with 64 GB memory and an Intel Core i7-9700 CPU, using R version 4.4.1 with machine epsilon 2.22×10^−16^.

### 3.1 Simulation Studies

#### 3.1.1 Case 1: Small Quadratic Forms with Algorithm 1

##### Setting 1: Sum of Central Chi-squares

Using the fact that a central chi-squared variable with *m* DFs can be expressed asm 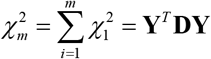, where **Y** follows *N*_*m*_ (**0, I**_*m*_) and **D** = diag(1,…,1) is a diagonal matrix with *m* ones, we evaluate the accuracy of Algorithm 1 for estimating *p*-values of different orders of magnitude by varying *q* and *m*, and compare the results with the exact *p*-values, 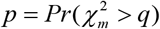, which can be accurately computed using the R base function *pchisq*. The following methods are also included in our comparisons: Liu-Tang-Zhang’s (LTZ’s, which approximate the distribution of the quadratic form using a non-central chi-squared distribution (Liu et al., 2009)), Davies’, Farebrother’s, Imhof’s methods (these four methods are implemented in the R package *CompQuadForm* (Duchesne and De Micheaux, 2010)), and the saddlepoint approximation approach (implemented in the R package *bigQF* (Lumley et al., 2018)). For each method, the absolute value of relative error (*ARE*) is used to assess the error, which is computed as 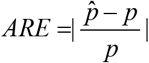, where 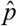 is the estimated *p*-value from each method and *p* is the exact *p*-value computed from *pchisq*. The numerical results are shown in Supplementary Table S1, and Fig. 1 presents a graphical comparison of different methods. Since LTZ’s method uses a scaled noncentral chi-squared variable to approximate the quadratic form *Q*, it obtains the exact form 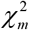 under this setting. Hence, its results are almost identical to the exact *p*-values, with *ARE*s close to the machine epsilon (see Supplementary Table S1, not shown in Fig. 1). Excluding LTZ’s method, Davies’ and Farebrother’s methods are more accurate than others for *p*-values larger than the machine epsilon, with Farebrother’s yielding slightly lower *ARE*s than Davies’. However, both fail for *p*-values close to or smaller than the machine epsilon, which is also observed by others (Chen and Lumley, 2019). Imhof’s fails when *p*-values are smaller than approximately 10^−8^ − 10^−9^. Saddlepoint approximation consistently performs well for *p*-values ranging from 10^−5^ to 10^−100^ with *ARE*s below 3%. MCMC-CE generally performs well for *p*-values ranging from 10^−5^ to 10^−100^ with *ARE*s below 10% when *m* ≤ 200, and it even slightly outperforms saddlepoint approximation when *m* = 5. However, its accuracy decreases when *m* increases to 250, and it converges very slowly with larger errors at higher dimensions (not shown here).

**Figure 1.**
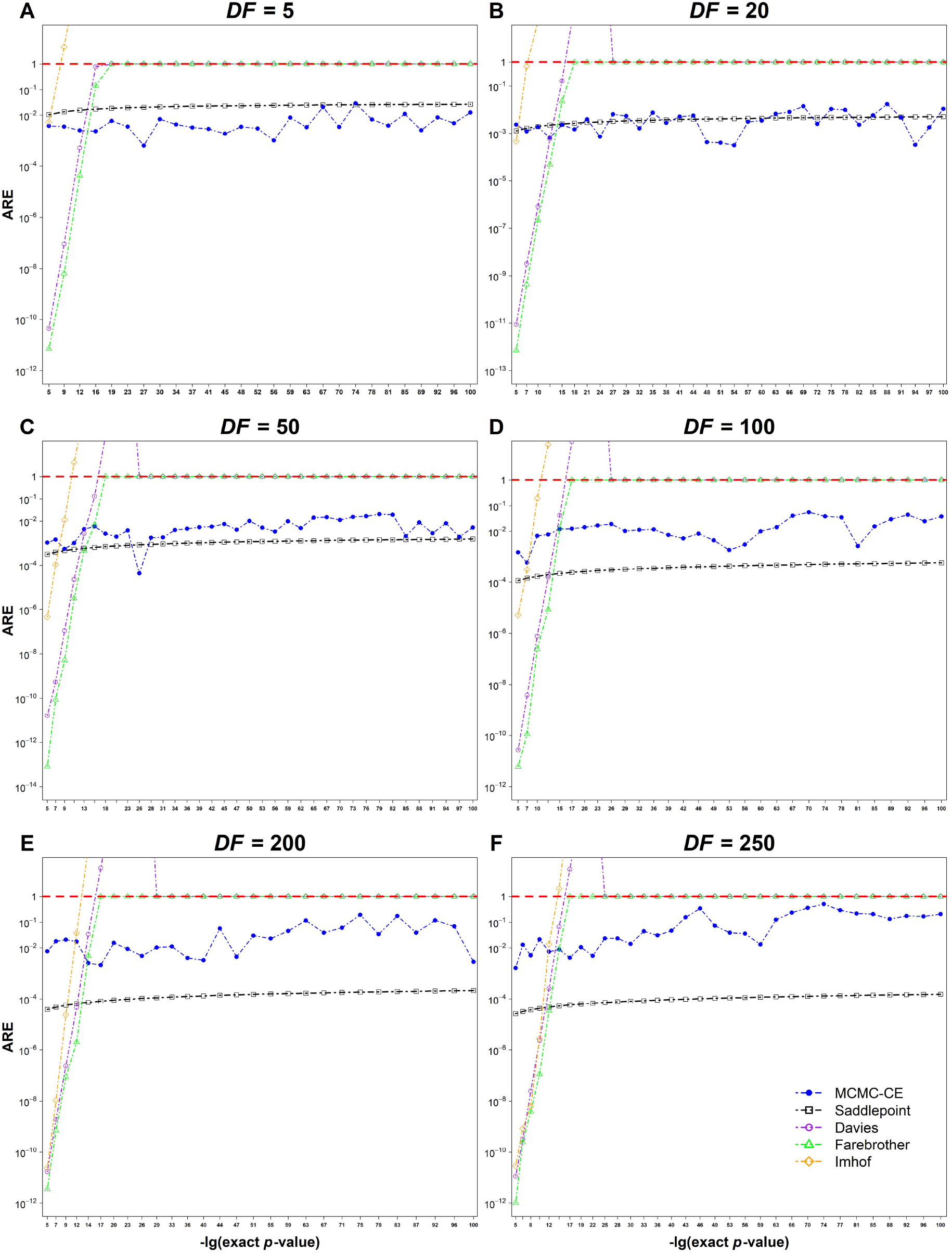
Comparisons between methods for estimating *p*-values associated with the sum of central chi-squared variables with different degrees of freedom (DFs). The details are described in Simulation Studies Setting 1, and the numerical results are presented in Supplementary Table S1. The metric plotted is the absolute value of the relative error (*ARE*) for each method. The dashed red horizontal line indicates an *ARE* of 100%.

##### Setting 2: Sum of Noncentral Chi-squares

Using the fact that a noncentral chi-squared variable with *m* DFs and NCP *δ* can be expressed as 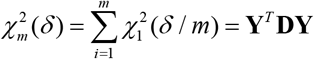, where **Y** follows 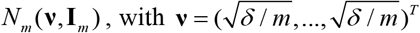, and **D** = diag(1,…,1) is a diagonal matrix with *m* ones, we evaluate the accuracy of Algorithm 1 similarly to Setting 1 by varying *m* and *δ*. The numerical results under this setting are shown in Supplementary Table S2, and Fig. 2 presents a graphical comparison of different methods. The performance of LTZ’s, Davies’, Farebrother’s and Imhof’s methods is similar to that in Setting 1. However, saddlepoint approximation is unreliable in this setting, with estimated *p*-values being quite liberal and several orders of magnitude smaller than the exact ones. MCMC-CE produces estimates within the same order of magnitude as the exact *p*-values for *p*-values down to approximately 10^−50^ for all tested values of DFs and NCPs, and its *ARE*s increase when DFs increase, NCP increases and *p*-values become smaller.

**Figure 2.**
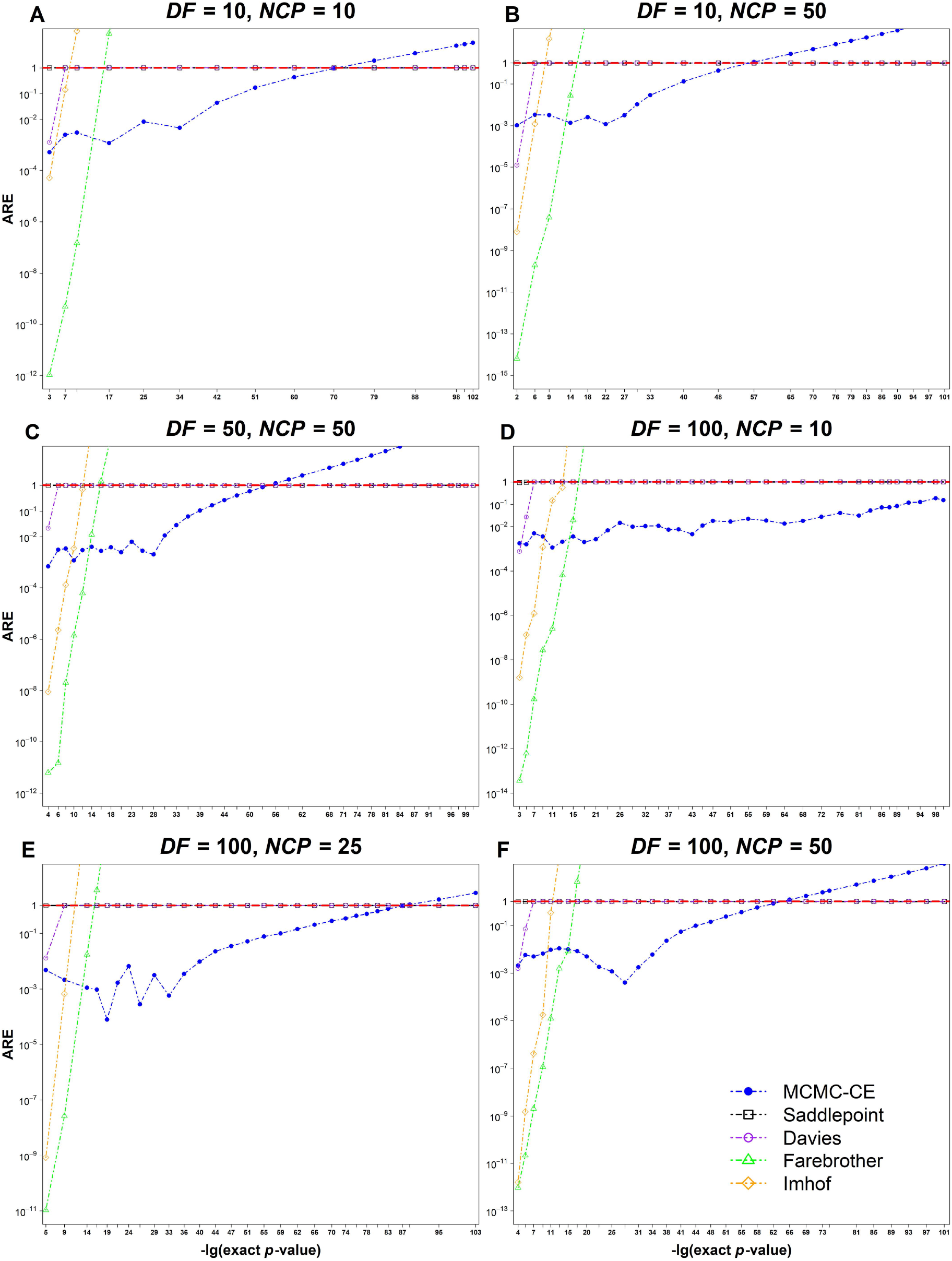
Comparisons between methods for estimating *p*-values associated with the sum of non-central chi-squared variables with different degrees of freedom (DFs) and different non-centrality parameters. The details are described in Simulation Studies Setting 2, and the numerical results are presented in Supplementary Table S2. The metric plotted is the absolute value of the relative error (*ARE*) for each method. The dashed red horizontal line indicates an *ARE* of 100%.

##### Setting 3: Low-dimensional Quadratic Forms

We compare different methods for estimating the *p*-values associated with two quadratic forms, *Q*_3_m and *Q*_7_, studied in (Duchesne and De Micheaux, 2010), which are defined as 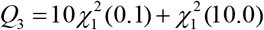 and 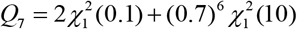. The exact *p*-values associated with *Q* and *Q* are unknown. Based on our simulations in the above settings and another study (Chen and Lumley, 2019), Farebrother’s method is the most accurate approach for small quadratic forms when their *p*-values are larger than the machine epsilon. Therefore, its results are used as the exact values for computing the *ARE*s of other methods when *p*-values are greater than 10^−15^. The results are shown in Supplementary Table S3, and Fig. 3 presents a graphical comparison of different methods’ *ARE*s for *p*-values > 10^−15^. Davies’ and Imhof’s methods yield similar results to those in Setting 1 and 2. MCMC-CE’s *ARE*s are controlled below ∼1% for *Q*_3_ and below ∼6% for *Q*_7_ for *p*-values > 10^−15^, and it still can provide a valid estimate for *p*-values < 10^−15^ when Farebrother’s, Davies’ and Imhof’s methods all fail. Saddlepoint approximation and LTZ’s are not accurate for *p*-values < 10^−5^ with very liberal estimates and large *ARE*s.

**Figure 3.**
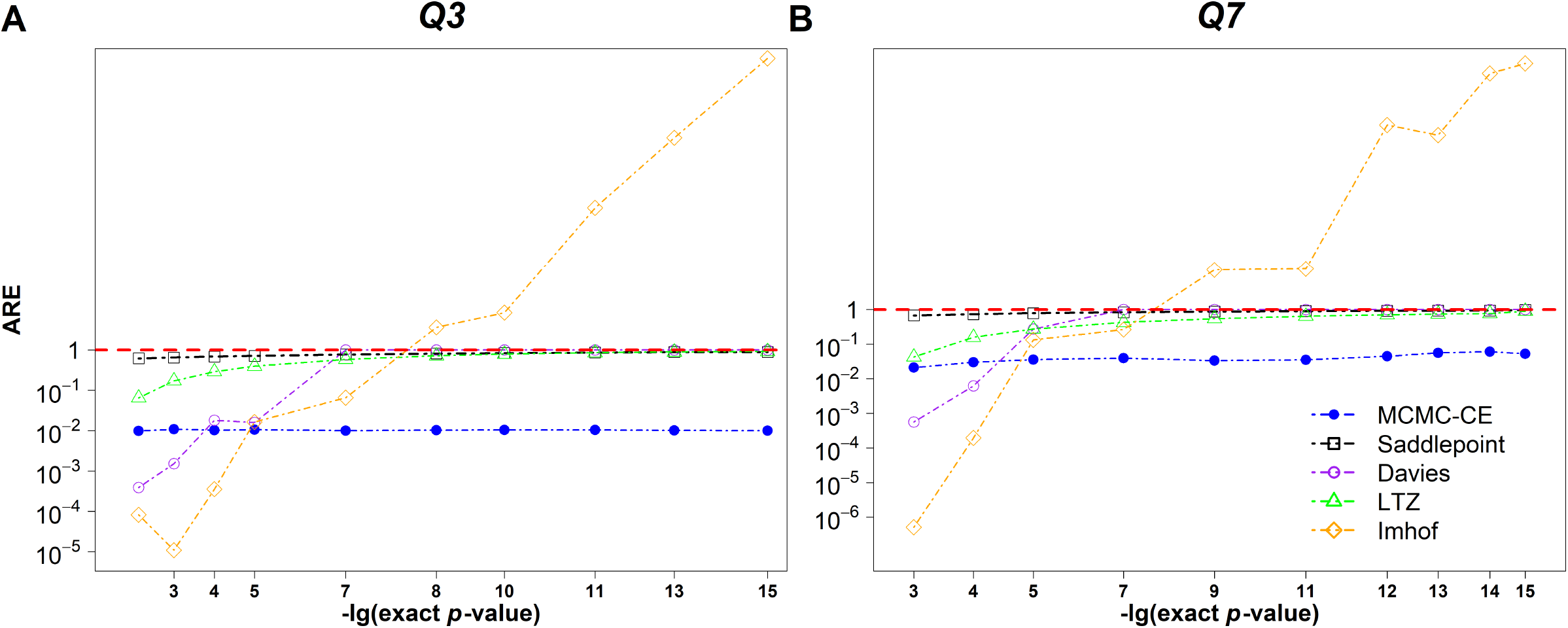
Comparisons between methods for estimating *p*-values associated with two quadratic forms *Q*_3_ and *Q*_7_. The details are described in Simulation Studies Setting 3, and the numerical results are presented in Supplementary Table S3. The metric plotted is the absolute value of the relative error (*ARE*) for each method. The dashed red horizontal line indicates an *ARE* of 100%.

#### 3.1.2 Case 2: Large Quadratic Forms with Algorithm 2

##### Setting 4: Runtime Comparison for Computing the Leading Eigenvalues

We first compare the computational time of trLanczos and SSVD for computing the *k* leading eigenvalues. In real-world data analysis, particularly in genomics, sometimes the matrix **A** associated with the quadratic form *Q* in (2) is available. However, in many other cases, the matrix **G, GG**^**T**^ = **A**, is available before **A**. In these cases, especially when **A** is large, directly working with **G** without computing **A** can greatly save computational time and memory, and procedures for computing the leading eigenvalues using trLanczos and SSVD when only **G** is available are proposed in (Lumley et al., 2018). We test the runtime of trLanczos and SSVD for computing different numbers of *k* leading eigenvalues with **A** available or only **G** available, with the rank *r* = 5,000 and 10,000, respectively. The results are shown in Table 1. The runtimes of the two methods are on the same order of magnitude, with SSVD faster when only **G** is available and *r* = 10,000. As expected, both methods are much faster than computing all eigenvalues using the *eigen* function in R base. These results are consistent with those reported in (Lumley et al., 2018).

**Table 1.** Runtime Comparisons of trLanczos and SSVD for Computing Different Number of Leading Eigenvalues^**a**^.

| $k$ | Method | $r = 5000$ | | $r = 10000$ | |
| --- | --- | --- | --- | --- | --- |
|  |  | A available | Only G | A available | Only G |
| 100 | trLanczos | 3.6 | 23.4 | 11.6 | 83.7 |
|  | SSVD | 5.9 | 20.6 | 14.8 | 50.3 |
| 150 | trLanczos | 5.5 | 26.7 | 15.3 | 107.2 |
|  | SSVD | 9.1 | 25.1 | 20.7 | 64.6 |
| 200 | trLanczos | 7.2 | 30.9 | 20.4 | 124.7 |
|  | SSVD | 12.3 | 28.4 | 25.9 | 80.6 |
| All ($k = r$) | R base *eigen*<sup>b</sup> | 37.3 | 125.3<sup>c</sup> | 218.6 | 650.4<sup>c</sup> |

**a.** Unit of time: second.
**b.** The runtime of computing all eigenvalues using R base *eigen* function is included for comparisons.
**c.** These times include the time of computing $\mathbf{A}$ as $\mathbf{A} = \mathbf{G}\mathbf{G}^T$ plus the time of computing all eigenvalues of $\mathbf{A}$ using the *eigen* function.

##### Setting 5: Large Quadratic Forms – Sum of Central Chi-squares

We compare different methods for estimating the *p*-values of large quadratic forms that can be written as sum of central chi-squares, corresponding to the forms of SKAT or *globaltest* test statistics under the null hypothesis of no association between the genomic features and the outcome. We find that Farebrother’s method, implemented in the R package *CompQuadForm*, is numerically unstable for large quadratic forms, suffering from underflow, an issue previously observed and explained in detail in (Chen and Lumley, 2019). Similarly, Imhof’s method, implemented in *CompQuadForm*, also exhibits numerical instability for large quadratic forms, and sometimes fails completely when the *p*-value is around 10^−5^. Therefore, these two methods are not included in our comparison. The quadratic forms used here are *Q*_1_ to *Q*_6_, with ranks ranging from 305 to 11,259, as given in (Chen and Lumley, 2019), which is simulated using the Markov Coalescent Simulator (Chen et al., 2009). Davies’, LTZ’s, saddlepoint approximation and MCMC-CE (Algorithm 2 with *k* = 150) methods are used to estimate the *p*-values associated with *Q*_1_ to *Q*_6_. The exact *p*-values associated with *Q*_1_ to *Q*_6_ are unknown. According to our simulations and (Chen and Lumley, 2019), Davies’ method is the most accurate approach for large quadratic forms where *p*-values are larger than the machine epsilon. Hence, its results are used as the exact values for computing the *ARE*s of other methods for *p*-values > 10^−15^. The results are shown in Supplementary Table S4, and Fig. 4 presents a graphical comparison of different methods’ *ARE*s for *p*-values > 10^−15^. LTZ’s estimates become highly liberal for *p*-values < 10^−3^, while Saddlepoint approximation’s *ARE*s remain controlled below 20% in all cases for *p*-values > 10^−15^. These results are consistent with the observations in (Chen and Lumley, 2019). MCMC-CE’s *ARE*s are slightly higher than those of the saddlepoint approximation, particularly when the rank of *Q* increases. However, they are controlled below around 25% for *p*-values > 10^−15^. MCMC-CE and saddlepoint approximation yield estimates of the same order of magnitude for *p*-values < 10^−15^.

**Figure 4.**
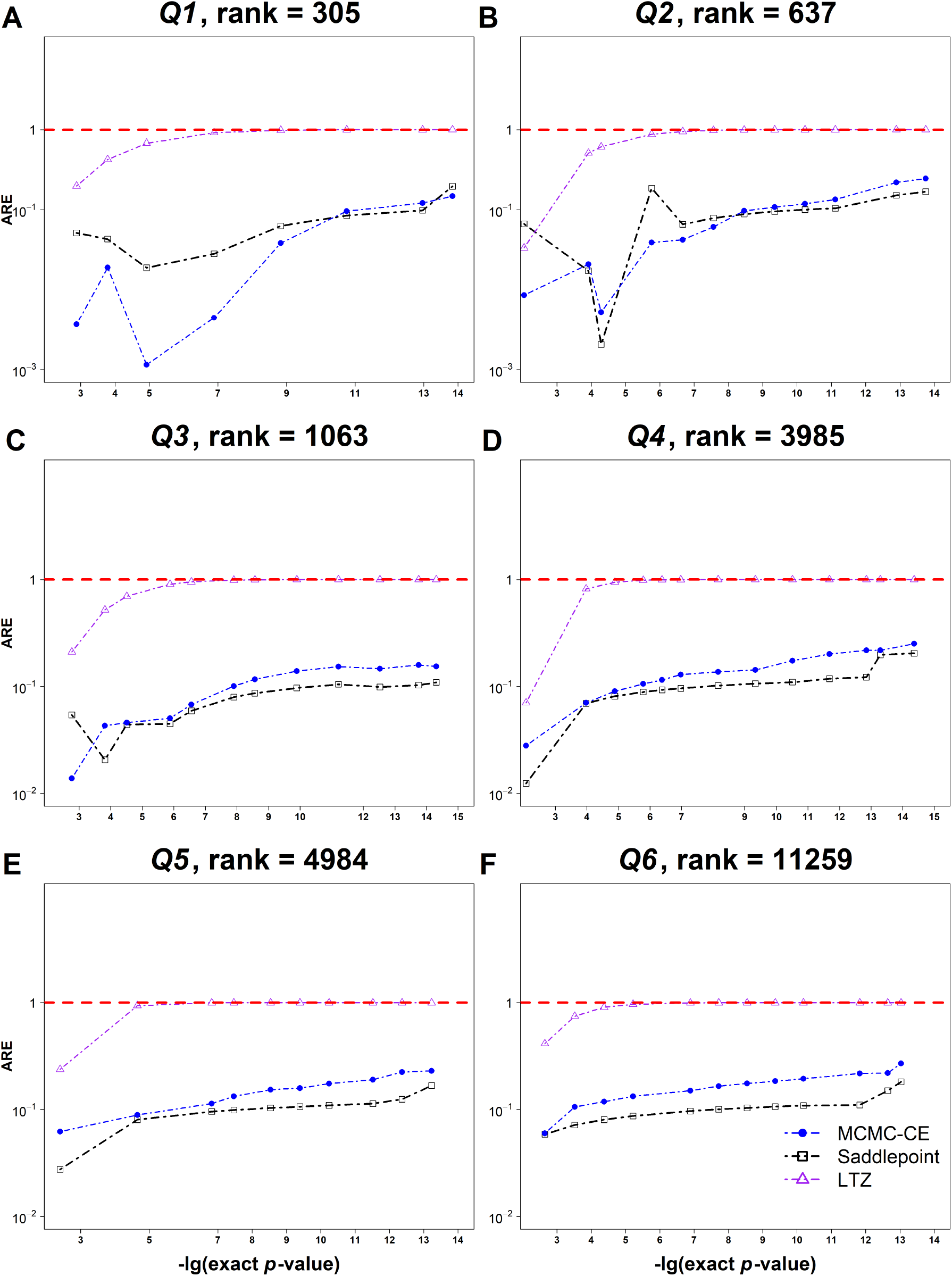
Comparisons between methods for estimating *p*-values associated with six large quadratic forms *Q*_1_ – *Q*_6_. The details are described in Simulation Studies Setting 5, and the numerical results are presented in Supplementary Table S4. The metric plotted is the absolute value of the relative error (*ARE*) for each method. The dashed red horizontal line indicates an *ARE* of 100%.

##### Setting 6: Large Quadratic Forms – Sum of Noncentral and Central Chi-squares

We evaluate different methods for large quadratic forms that consist of the sums of both noncentral and central chi-squares. We assume that approximately the first 1% to 5% of the terms in (5) are noncentral chi-squares. Particularly, the quadratic forms used here are generated from *Q*_1_ to *Q*_6_ in Setting 5 and take the following form,

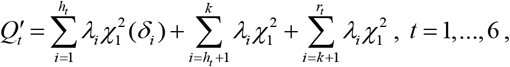

where *t* is the index denoting *Q*_*t*_′ is generated from which of *Q*_1_ to *Q*_6_, *λ*_*i*_’s, *i* = 1,…, *r*_*t*_, are the *r*_*t*_ ranked eigenvalues of *Q*_*t*_, *k* is the number of leading eigenvalues used in Algorithm 2, which equals150, *h*_*t*_ is the number of non-central chi-squared terms, which is specified as *h*_*t*_ = _└_*ρ*_*t*_ *r*_*t* ┘_, with _└_·_┘_ the floor function, *ρ*_1_ = 5%, *ρ*_2_ = *ρ*_3_ = 3%, *ρ*_4_ = *ρ*_5_ = 2%, *ρ*_6_ = 1% and the NCPs *δ*_*i*_’s, *i* = 1,…, *h*_*t*_, randomly drawn from a uniform *U* (0, 5) distribution. This simulation setting represents a scenario in genomics where *ρ*_*t*_ genomic features among the *r*_*t*_ features are associated with the outcome variable.

The performance of LTZ’s, Davies’, saddlepoint approximation and MCMC-CE (Algorithm 2 with *k* = 150) is evaluated for these newly generated quadratic forms *Q*_1_′ to *Q*_6_′. Similar to Setting 5, Davies’ results are used as the exact *p*-values for computing the *ARE*s of other methods for *p*-values > 10^−15^. The results are shown in Supplementary Table S5, and Fig. 5 presents a graphical comparison of different methods’ *ARE*s for *p*-values > 10^−15^. Saddlepoint approximation’s and LTZ’s estimates become highly liberal when *p*-values drop below 10^−5^. MCMC-CE’s *ARE*s increase as the rank of *Q* increases, but are controlled below around 35% for *p*-values > 10^−15^ in all cases. For *p*-values < 10^−15^, saddlepoint approximation’s and LTZ’s estimates are several orders of magnitude smaller than those of MCMC-CE, as also observed in Setting 3.

**Figure 5.**
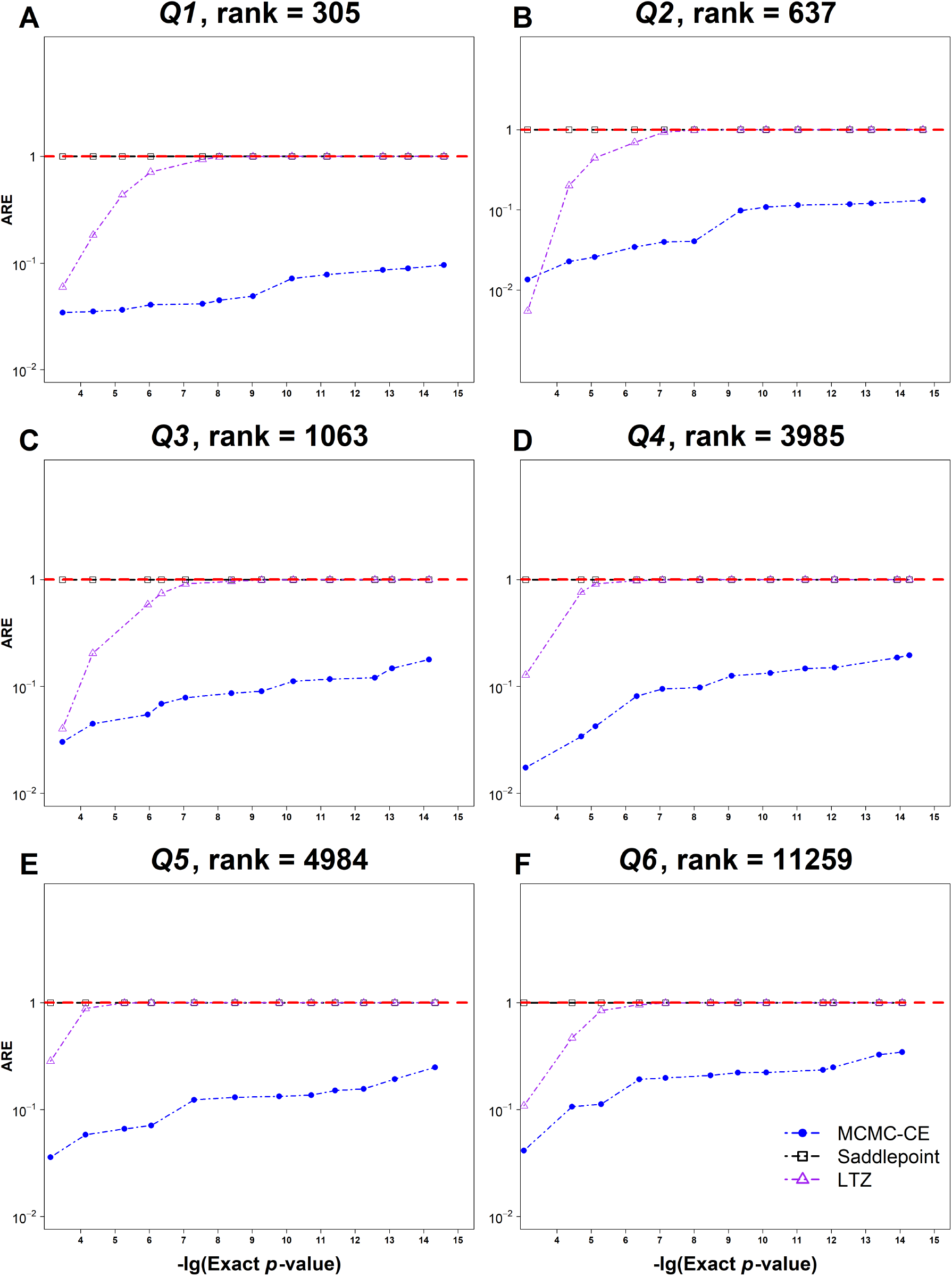
Comparisons between methods for estimating *p*-values associated with six large quadratic forms *Q*_1_′ – *Q*_6_′. The details are described in Simulation Studies Setting 6, and the numerical results are presented in Supplementary Table S5. The metric plotted is the absolute value of the relative error (*ARE*) for each method. The dashed red horizontal line indicates an *ARE* of 100%.

### 3.2. Application to Genomics Data

#### 3.2.1. Application to a GWAS Dataset Using SKAT

We demonstrate the strength of MCMC-CE for computing small *p*-values for both small and large quadratic forms by a mouse GWAS study using SKAT (Wu et al., 2010, Wu et al., 2011). Ideally, SKAT with large quadratic forms is often applied to large-scale human GWAS study (e.g. with over 10,000 participants and over one million SNPs (Lumley et al., 2018)). However, the individual-level genotype and phenotype data from such large-scale human GWAS are usually not publicly available due to privacy concerns. Therefore, we use a publicly available small-scale mouse GWAS dataset here. The goal of our analysis below is to demonstrate the computational efficiency and strength of MCMC-CE for computing extremely small *p*-values with the quadratic form statistics such as the one used in SKAT, and it should be noted that SKAT with large quadratic forms may not be suitable to be applied to this dataset. This dataset is generated from a population of around 2,000 heterogeneous stock mice with over 100 traits (Valdar et al., 2006). The outcome variable is the serum concentration of high-density lipoprotein (HDL), and the goal is to identify the genomic locations (group of adjacent SNPs) on the chromosomes significantly associated with HDL levels using the SKAT test. After pre-processing, the dataset contains 1,640 subjects and 10,990 SNPs with complete HDL and genotype data. We apply a sliding window approach with window sizes of 50, 500, and 5000 SNPs respectively to scan the whole genome (e.g. for a window size of 50 SNPs, adjacent SNPs are grouped into sets of 50, and the SKAT is applied to each set. The test continues across the entire genome. This type of genome scan is widely used in GWAS (Wu et al., 2011)). For a window size of 50, Algorithm 1 is applied using all the eigenvalues of the SKAT statistic. For window sizes of 500 and 5000, Algorithm 2 is applied with *k*, the number of leading eigenvalues used, equals 150. Table 2 shows the computational times and smallest *p*-values estimated for different window sizes. These results demonstrate that MCMC-CE is capable of computing extremely small *p*-values for large quadratic forms within a reasonable and affordable timeframe on a desktop PC.

**Table 2.** Runtime and Minimum *P*-values Estimated by MCMC-CE under Different Sliding Window Sizes for the Mouse GWAS Data.

| Window Size | No. Windows Tested | Minimum $P$-value | Time |
| --- | --- | --- | --- |
| 50 | 4356 | $2.97 \times 10^{-71}$ | 1.13 h |
| 500 | 10491 | $3.80 \times 10^{-62}$ | 5.23 h |
| 5000 | 5991 | $2.27 \times 10^{-81}$ | 15.63 h |

#### 3.2.2. Application to an RNA-Seq Dataset for Pathway Enrichment Analysis

We apply MCMC-CE to an RNA-Seq gene expression dataset from the Cancer Genome Atlas breast cancer (TCGA-BRCA) project (Cancer Genome Atlas, 2012) for pathway enrichment analysis using *globaltest*. After pre-processing and excluding the male patient samples due to their small sample size (*n* = 14), this dataset includes gene expression levels measured in fragments per kilobase per million mapped fragments (FPKMs) of 24,622 genes from 1,210 female breast cancer patient samples, of which 1,098 are primary breast tumors and 112 are normal breast tissues. We apply the model and test implemented in the R package *globaltest* (Goeman et al., 2004) to analyze the 361 pathways curated in the Kyoto Encyclopedia of Genes and Genomes (KEGG) database (Kanehisa and Goto, 2000). Specifically, a logistic regression model is fitted with tumor or normal status as the outcome variable and race, age and expression levels of the genes in the pathway of interest as predictors, and the quadratic form of the *globaltest* statistic is then used to test the significance of the association between the genes in the pathway and the outcome. Since our goal is to demonstrate the strength of MCMC-CE for estimating small *p*-values of quadratic forms, the following screening test is applied to filter out less-significant pathways with large *p*-values: the approximated *p*-values are first computed for all pathways using the scaled chi-squared distribution implemented in *globaltest*, and 67 pathways with the approximated *p*-values less than 10^−5^ are selected. MCMC-CE is then used to accurately estimate the *p*-values associated with these 67 significant pathways. The results are shown in Supplementary Table S6, and Table 3 presents the *p*-values for the top 10 most significant pathways. MCMC-CE is capable of estimating small *p*-values down to the order of magnitude of 10^−65^. For comparisons, Farebrother’s and Davies’ methods are also applied to these 67 pathways. As expected, both of them fail to provide valid estimates for *p*-values smaller than the machine epsilon. For those *p*-values between 10^−15^ and 10^−5^, the results of Farebrother’s, Davies’ and MCMC-CE are very similar.

**Table 3.**
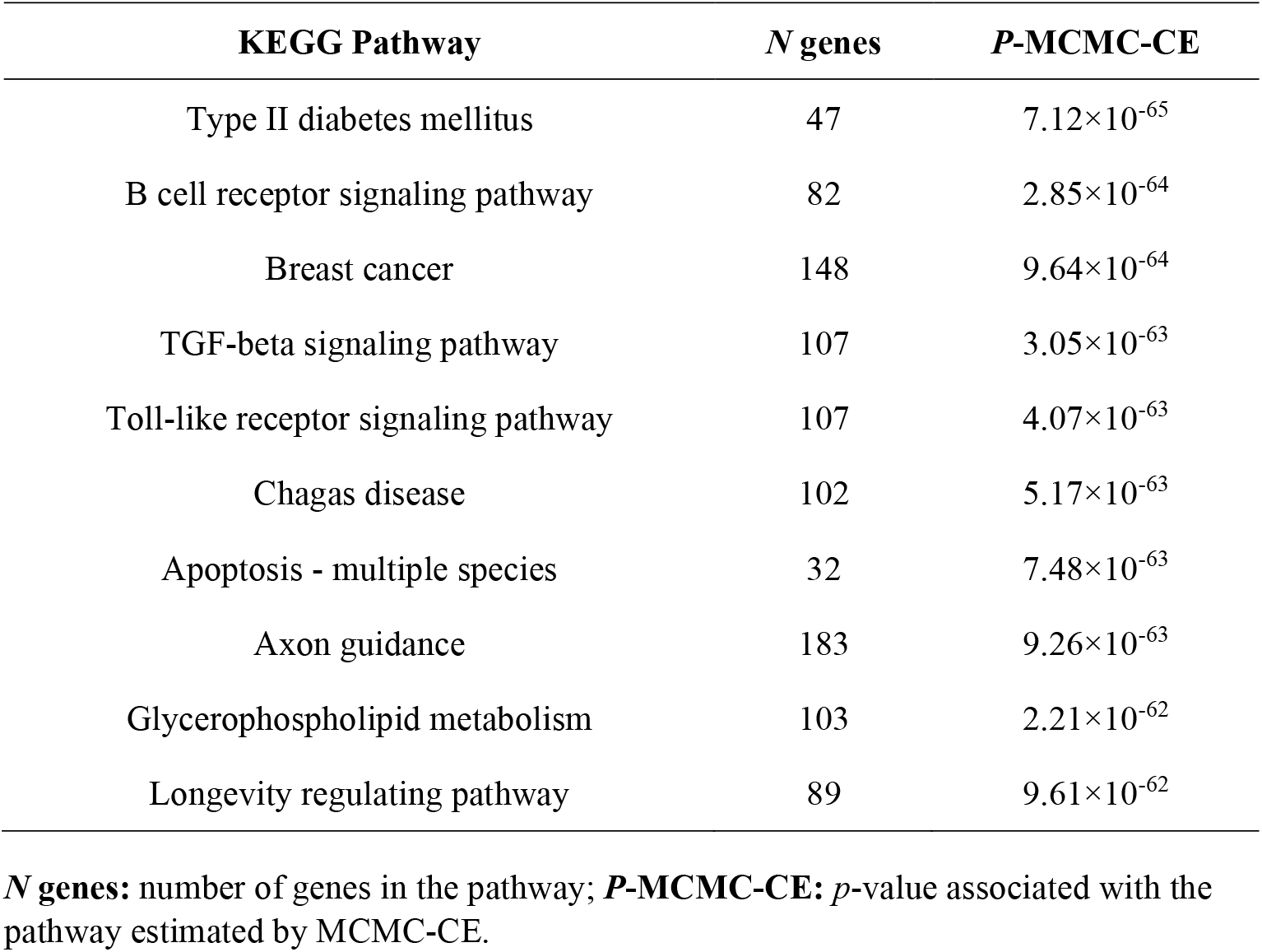
Top 10 KEGG Pathways Significantly Associated with Breast Cancer Ranked by Their *P*-values Estimated by MCMC-CE in TCGA-BRCA RNA-Seq Dataset.

## 4. CONCLUSION AND DISCUSSIONS

In this work, we propose MCMC-CE, an efficient and scalable algorithm for accurately estimating small *p*-values of large-scale quadratic forms. Through extensive simulation studies and real-world genomic applications, we demonstrate that MCMC-CE provides advantages in numerical accuracy and computational reliability compared to existing methods, including Davies’, Imhof’s, Farebrother’s, LTZ’s and the saddlepoint approximation methods. Our approach overcomes numerical challenges encountered in these traditional methods, making it a practical and robust solution for computing extremely small *p*-values in large-scale genomic studies using quadratic forms.

Despite its strengths, our simulation studies indicate that the accuracy of MCMC-CE for large quadratic forms decreases as the dimension of the quadratic form increases and *p*-values become smaller, as reflected by its increased *ARE*s. This is expected because, as the number of the remainder terms in (5) increases, the approximation of their sum using a single scaled noncentral chi-squared variable (as in Algorithm 2) becomes less precise. To further improve accuracy, alternative approximation schemes can be considered. One possible approach is to approximate the sum 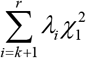 in (5) with another scaled noncentral chi-squared variable, 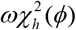, with three parameters *ω, ϕ* and *h*, which should provide a better approximation than 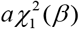 in principle. That is, approximate the distribution of *Q* in (5) using the distribution of

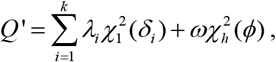

where the three parameters of 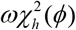 can be determined by moment matching with 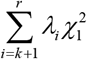. However, computing these parameters is non-trivial and presents additional computational challenges. Furthermore, MCMC-CE requires that the dimension of *Q* ‘, *k* + *h*, does not exceed 250 in practice, meaning that the choice of *h* must be constrained to ensure computational feasibility, which further complicates this approximation scheme. We consider exploring such alternative approximation techniques that improve the accuracy of MCMC-CE for high-dimensional quadratic forms as future work.

## Acknowledgements

This work was partially from V.Q.O.’s dissertation while he was a doctoral student at Augusta University, and Drs. Jie Chen and Dustin Pluta at Augusta University have critically read his dissertation and provided helpful discussions.

## Data Availability Statement

The mouse GWAS dataset is publicly available at https://wp.cs.ucl.ac.uk/outbredmice/heterogeneous-stock-mice. The RNA-Seq gene expression dataset from TCGA-BRCA project is publicly available at https://portal.gdc.cancer.gov/projects/TCGA-BRCA. R code for implementing the algorithms and performing the analyses is available at https://github.com/shilab2017/MCMC-CE-quadratic-forms.

